# On the joint effect of endogenous spatial attention and defocus blur on acuity: attentional limit to the resolving power of the eye

**DOI:** 10.1101/2020.08.25.266700

**Authors:** E De Lestrange-Anginieur, TW Leung, CS Kee

## Abstract

Defocus blur and spatial attention both act on our ability to see clearly over time. However, it is currently unknown how these two factors interact because studies on acuity resolution only focused on the separate effects of attention and defocus blurs. In this study, resolution acuity was measured along the diagonal 135°/315° with horizontal, at 8° eccentricity for clear and blur Landolt C images under various manipulations of covert endogenous attention. We observe that attention not just improves the resolution of clear stimuli, but also modulates the resolution of defocused stimuli for compensating the loss of resolution caused by retinal blur. Our results show, however, that as the degree of attention decreases, the differences between clear and blurred images largely diminish, thus limiting the benefit of an image quality enhancement. It also appeared that attention tends to enhance the resolution of clear images more than blurred targets, suggesting potential variations in the gain of vision correction with the level of attention. This demonstrates that the interaction between spatial attention and focus plays a role in the way we see things. In view of these findings, the development of adaptive (neuro-optical) interventions, which adjust the eye’s focus to attention, may hold promise.

**Significance statement:** Visual technologies are now attaining a degree of extreme sophistication and diversity, which allows more comprehensive, but often complex manipulations of the optical image formed onto the back of the eye. It is therefore an enigma how those fine and immersive manipulations of the sensory environment are integrated in the brain. In this study, we show that the resolving power of the eye can depend complexly on the interaction between spatial attention and focus. This discovery suggests that perception might be advantageously guided by technologies tailoring optical focus to individual attentional patterns.

**Graphical abstract:** 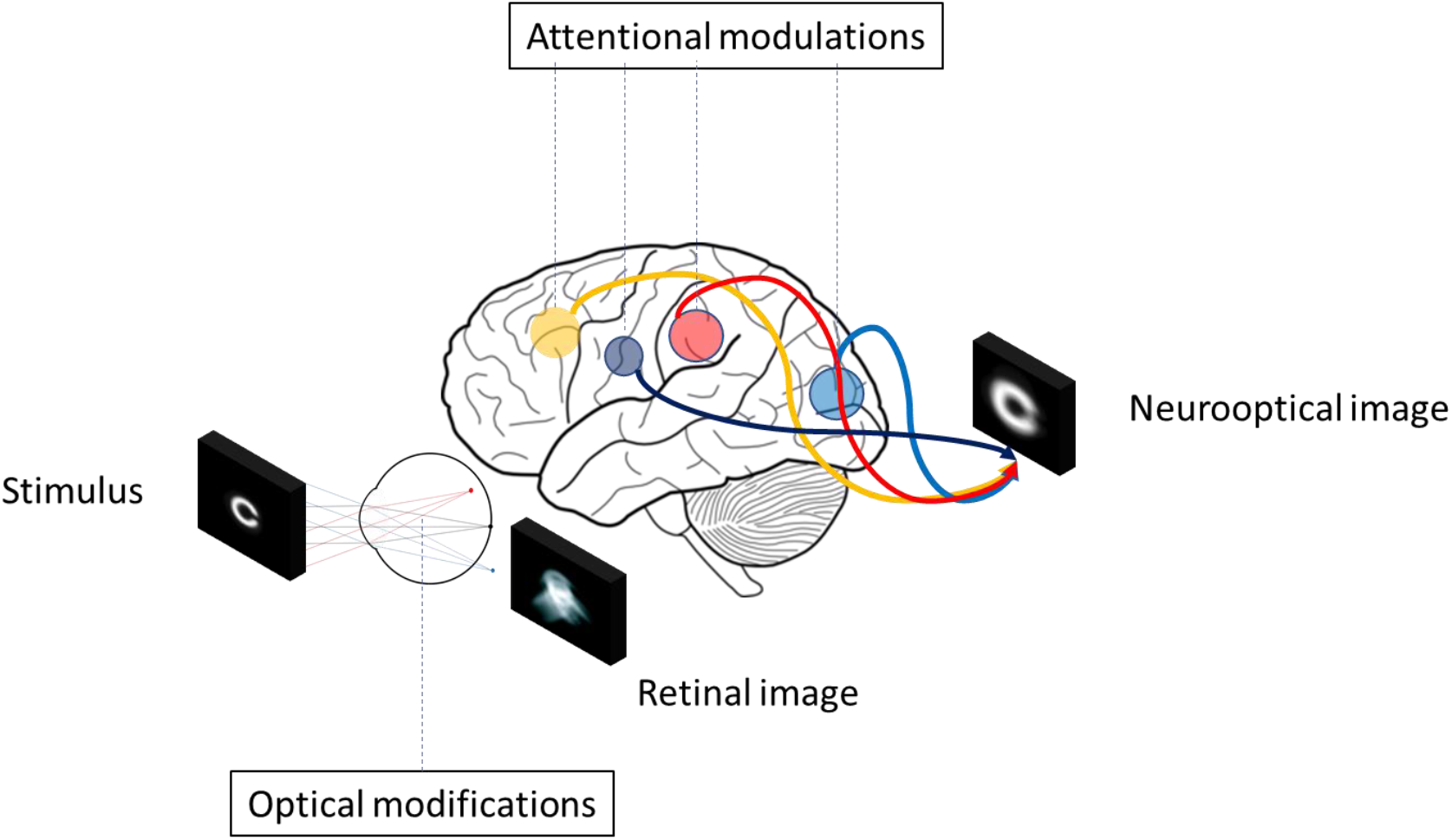

Proposed neuro-optical mechanism of transformation of stimulus by spatial attention and blur.

---

Our ability to perceive the constituents of a visual stimulation strongly depends on the optical focus of the eye that defines the state for which the smallest spatial feature of the environment can be resolved. The control of optical focus has been one the most significant challenges of optics, and continues to drive a large number of visual technologies, such as ophthalmic visual aids (such as spectacles, contact lenses), but also display technologies (Virtual reality, vision-correcting light field display) aimed at improving and correcting the human sensory experience. Often controlling more than one visual target, those technologies have become both adaptive [**1-2**] and more complex [**3**], capable of manipulating image quality over time and space, which may broaden our experience and interaction with the environment. However, even with such heightened optical manipulation, predicting the impacts of an optical correction on visual function remains challenging due to an uncertainty as to how and to what extent optical modification affects the way retinal images are processed by the brain. Considering a capacity-limited visual system [**4**] and the infinite sum of information present in the visual world, augmentation of optical information could be traded off by increase amount and time for neural computation to access a neural representation, veridical with respect to the retinal image. It is known that usually not all information entering the eye is effectively processed over our visual field, with some spatial and temporal information never reaching consciousness or receiving attention [**5**]. Incompleteness of processing or absence of attention could limit the gain of correcting or augmenting optical signals. A vivid illustration of this phenomenon, the inattentional blindness paradigm [**6**], is the failure to report a simple, suprathreshold stimulus or a stimulus attribute of the visual field in the absence of attention, when the eye is engaged in another attention demanding primary task. Despite this deficit, it has been shown that some amount of processing still occurs in absence of attention, when the stimulus fails to be detected [**7-8**]. It has been demonstrated that the effect of attention strongly depends on the type of spatial information [**9**], and so our visual experience may vary with the characteristics of the environment. However, the effect of movement of attention on the processing of optical signals (i.e., signals having a complex distribution of spatial frequency - dependent contrast) has received little attention. Thus, to date, it is unknown to what extent attention contributes to the perception of the fine optical signals unveiled by ocular correction. A recent examination of the relationship between blur and attention proposed that blur detection might be pre-attentively processed by the visual system [**10**]. Interestingly, this suggests that, even in the absence of attention, when a stimulus attribute failed to be reported, visual blur might be partly processed. Nevertheless, the degree to which blur analysis remains independent from attention deployments is still unclear. Comparing the effect of blurred and clear images under variation of attention could help determine the potential benefit of correcting visual blur under real life contexts, when attention varies. To comprehensively examine this relationship, several forms of attention and type of blur may be of interest. This study focused on the interaction between endogenous covert attention and visual blur.

In covert attention [**11**], sensory enhancement occurs, without eye movements, by granting priority of processing to a certain location of the visual field, which usually takes place at the expense of processes in other regions stimulating the retina. This prioritization can occur via two modes of control: The first mode, exogenous attention, is a rapid, reflex, automatic deployment of attention in response to external visual stimulation, such as an abrupt, peripheral visual event, requiring an immediate response. The second mode, endogenous attention, is under conscious, voluntary control and can be directed according to the observer’s visual goal provided there is sufficient time of activation. In real life, both endogenous and exogenous factors [**12**] affect the deployment of attention, which implies a complex interaction between environmental characteristics, neural architecture, and cognitive behaviors. A major interest of endogenous attention is that, unlike exogenous attention, it can be flexibly sustained at a position of the visual field suitable for the task requirements [**13-15**]. In laboratory conditions, the strength of endogenous attention can be manipulated via a central symbolic cue (e.g., an arrow pointing towards the region of the cued location), and takes about 300ms from cue onset to deploy.

Neurophysiological studies have shown that as attention deploys, sensitivity (contrast gain [**16**]) or/and firing rate (response gain [**17**]) of neurons are increased, altering the relationship between stimulus contrast and neurons response. This can manifest in an enhancement of several perceptual tasks and stimulus properties, including resolution acuity [**18-27**], resolvable and apparent contrast [**28-29**]. Failure to adequately direct attention to environmental stimulation has been shown to drastically impair sensory performance. How does this attentional variability affect an optical ocular correction? Recent behavioral studies demonstrate the existence of a preferential enhancement of neurons tuned to high spatial frequency [**30**], with endogenous attentional modulation at both low and high spatial frequencies [**31**]. This differential effect suggests that variations of neural filtering by the focus of attention could potentially alter the *complex* appearance of broadband stimuli, having more than one spatial frequency. Such alteration could take place under various forms depending on the spatial extent of attention modulation in broadband images containing more than one spatial frequency. It is however unknown how attention varies with the complex distribution of spatial frequency pattern set by retinal blur. An account of both optical transfer function and neural transfer function could predict the neurooptical transformation of individual retinal images [**32**]. Unfortunately, because previous measurements of the contrast sensitivity function [**31, 33**] did not control the effect of blur on attention, it remains unknown how attention reshapes the neural transfer function, which is the neural part determining perception independently of the optics of the eye. One of the questions that ensues is whether the modulation of attended and unattended stimulus is affected by the differentiation of ocular optical filtering across spatial frequencies and the variation of ocular blur across individuals. For example, the attenuation of high spatial frequencies in uncorrected stimuli could increase or decrease focus of attention with neural adaptation to the image the visual system decodes.

In this study, we were interested to examine whether, or not, there is a possible interaction between blur and attention on acuity, and its potential implication when correcting the eye via optical or attentional manipulations. We noted that previous studies assessing the impact of blur in acuity [**34-40**] systematically overlooked the effect of attention on acuity, and vice versa. To the best of our knowledge, this is the first studies that examined the joint effect of blur and spatial attention on resolution acuity. Since attention can affect differentially spatial frequencies [**9,30-32**], the variation of the spatial frequency distribution with retinal blurs could vary the effect of attention, and thus have practical implications when correcting retinal blurs with an ophthalmic correction. Then, to which extent, and how? Here, we hypothesized that blurred stimuli, exhibiting greater attenuation of contrast-dependent spatial frequency [**41**], may benefit less from attentional enhancement, as compared to clear stimuli. To test the effect of retinal defocus blur on attention, we used the “source method” proposed by Haig et al [**42**], which is a widely used method for assessing the visual impact of ocular aberrations [**34-40**], and the most optimum approach for simulating spatiotemporally-varying blurs stimuli, which cannot be simulated with adaptive optics. This study shows evidence that the deployments of attention in non-foveal locations act on the visual effect of correction by altering the acuity distinctions between clear and blur stimuli. These findings demonstrate the existence of an attentional modulation of the effect of retinal blurs, and may motivate the development of novel interventions based on the adjustment of focus.

## Results

Adapting a Posner’s cueing paradigm introduced by other researchers [**18**], resolution acuity was tested under the manipulation of blur and covert endogenous spatial attention. The visual stimuli consisted of a pair of Landolt C letters (Fig. 1, step 3; “target” and “non-target”, with different gap orientations in each) briefly displayed at two locations in the near-periphery of the visual field at 8° eccentricity. Spatial attention was manipulated using three cueing conditions (Fig. 1, step 2), whereby a cue preceded the Landolt C stimuli: (1) cued condition, in which a cue was displayed and pointed at the upcoming target Landolt C location; (2) uncued condition, in which the cue pointed at the non-target upcoming Landolt C location; and (3) neutral condition, in which two cues were simultaneously displayed and pointed at both target and non-target locations. In cued and uncued trials, observers were required to attend to a cued location (either north-west [NW] or south-east [SE]) while fixating on a small central cross. In neutral trials, observers were required to spread their attention and focus on both locations (i.e., NW and SE). At the end of the stimulus presentation sequence, the location of the target Landolt C (Fig. 1, step 4) was indicated by a response cue, which consists of a central line symbol pointing towards one of the two stimuli.

**Figure 1.**
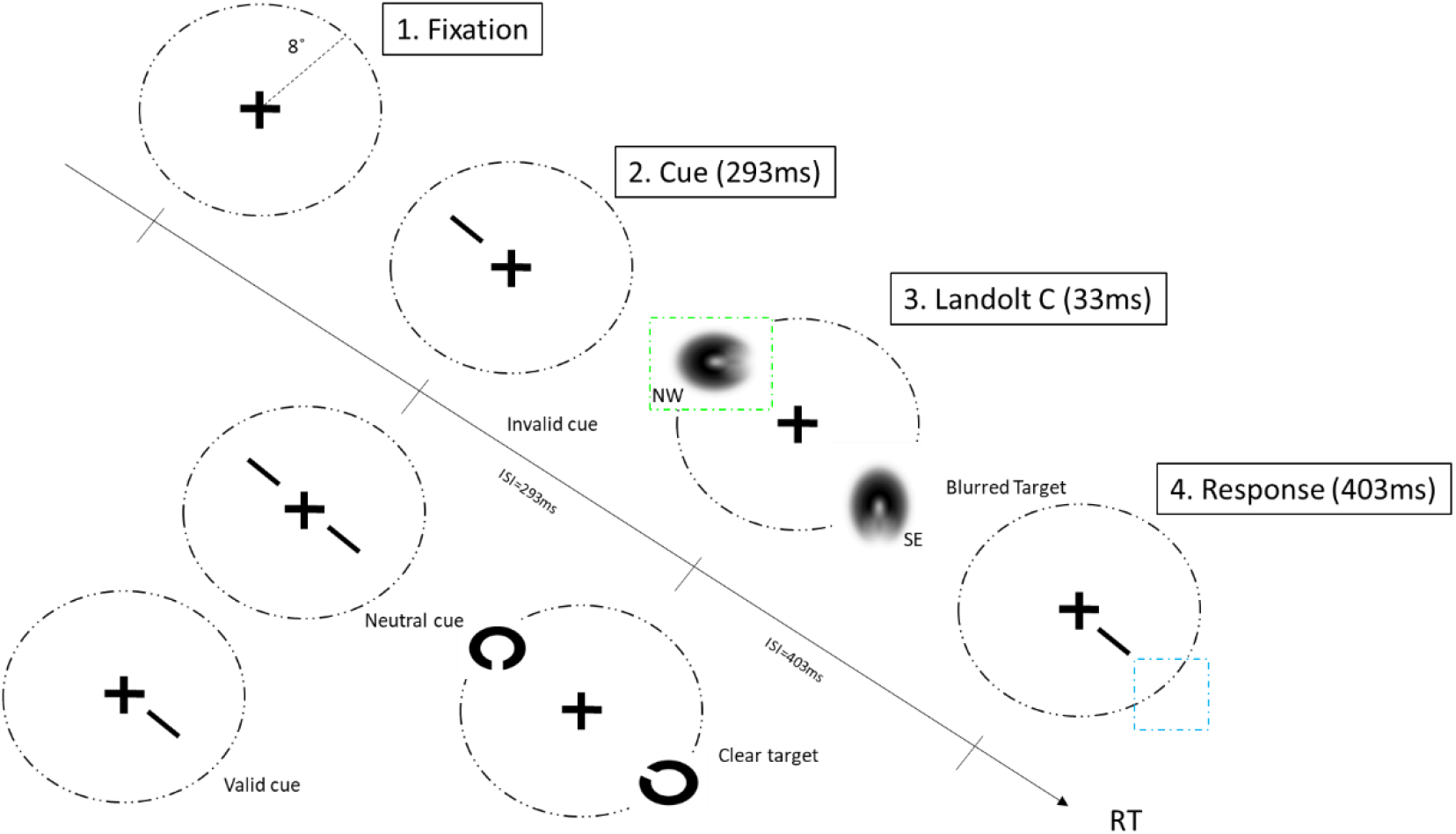
Visual acuity in the near-peripheral retina. In this example, the subject was required to respond to the Landolt C located in the SE quadrant. Visual acuity was tested at two locations (NW and SE) under various blur levels (blurred and clear) and cueing conditions (cued, uncued, and neutral). NW, north-west; SE, south-east; RT, response time.

The visual task was to identify the orientation of the gap in the “target” Landolt C (6AFC; see Methods for details), as indicated by the response cue. Visual acuity performance was measured under both clear and blurred conditions (Fig. 1, step 3). The gap size was controlled using an adaptive 1-down-1-up staircase procedure. A computerized image processing technique was employed to simulate the effects of a moderate amount of optical defocus on the retinal image of the Landolt C stimulus [**43**] (defocus blur, two waves of RMS wavefront error, about 1.25 diopters; see Methods for details). We chose to use a *clear* attentional cue and fixation stimuli in all the conditions so that the magnitude of the attention deployment was identical when comparing clear and blurred target conditions. There was a total of 12 interleaved staircases, for a total of 12 conditions (i.e., two retinal locations x three cue conditions x two blur levels). In addition to visual acuity (VA), response time (RT) was also measured.

Eleven young adults with corrected-to-normal vision participated. The visual acuity and response time data were processed using a three-way repeated measures analysis of variance (RANOVA) test (retinal location: NW and SW; visual blur: clear and blurred; spatial cueing: cued, uncued, and neutral), and post-hoc pairwise comparisons were performed with the Bonferroni correction using SPSS.

### Spatial location

Spatial location of the target had a statistically significant effect on the overall performance for RT (three-way RANOVA: F(1,10)= 5.473 p=.041, *η*^2^=0.364), but not for VA (three-way RANOVA: F(1,10)=2.313 p=.159, *η*^2^=0.188). There was no significant interaction between spatial cueing effect and spatial location for either RT (three-way RANOVA RT: F(1.184,20)=3.227, p=.094, *η*^2^ =0.244) or VA (three-way RANOVA VA: F(2,20)=.081 p=.922, *η*^2^=0.008), suggesting that spatial cueing effect was little affected by the position of the target along the diagonal (135°/315° with horizontal).

### Spatial cueing

Spatial cueing was found to significantly influence VA (Fig. 2a, three-way RANOVA: F(1.109,11.091)=21.962, p=0.001, *η*^2^=0.687), as reported previously [**18-27**], and response time (Fig. 2c, three-way RANOVA: F(2,20)=28.919, p<0.001, *η*^2^ =0.743). Visual acuity performance was superior for cued stimuli when compared to that for neutral stimuli (Bonferroni post-hoc test VA: mean difference=1.981, arcmin, p=0.012; RT: mean difference=-0.052 ms, p=0.019) or uncued stimuli (Bonferroni post-hoc test VA: mean difference=5.963, arcmin, p=0.002; RT: mean difference=0.160 ms, p=0.001). Using the neutral condition as a reference, the data sets were replotted as changes in visual acuity in Fig. 2b and changes in response time in Fig. 2d. Note that positive values represent an increase in visual acuity performance, and vice versa. The overall cost of attending to an incorrect location was greater than the benefit of attending to a correct location by two-fold for both RT and VA.

**Figure 2.**
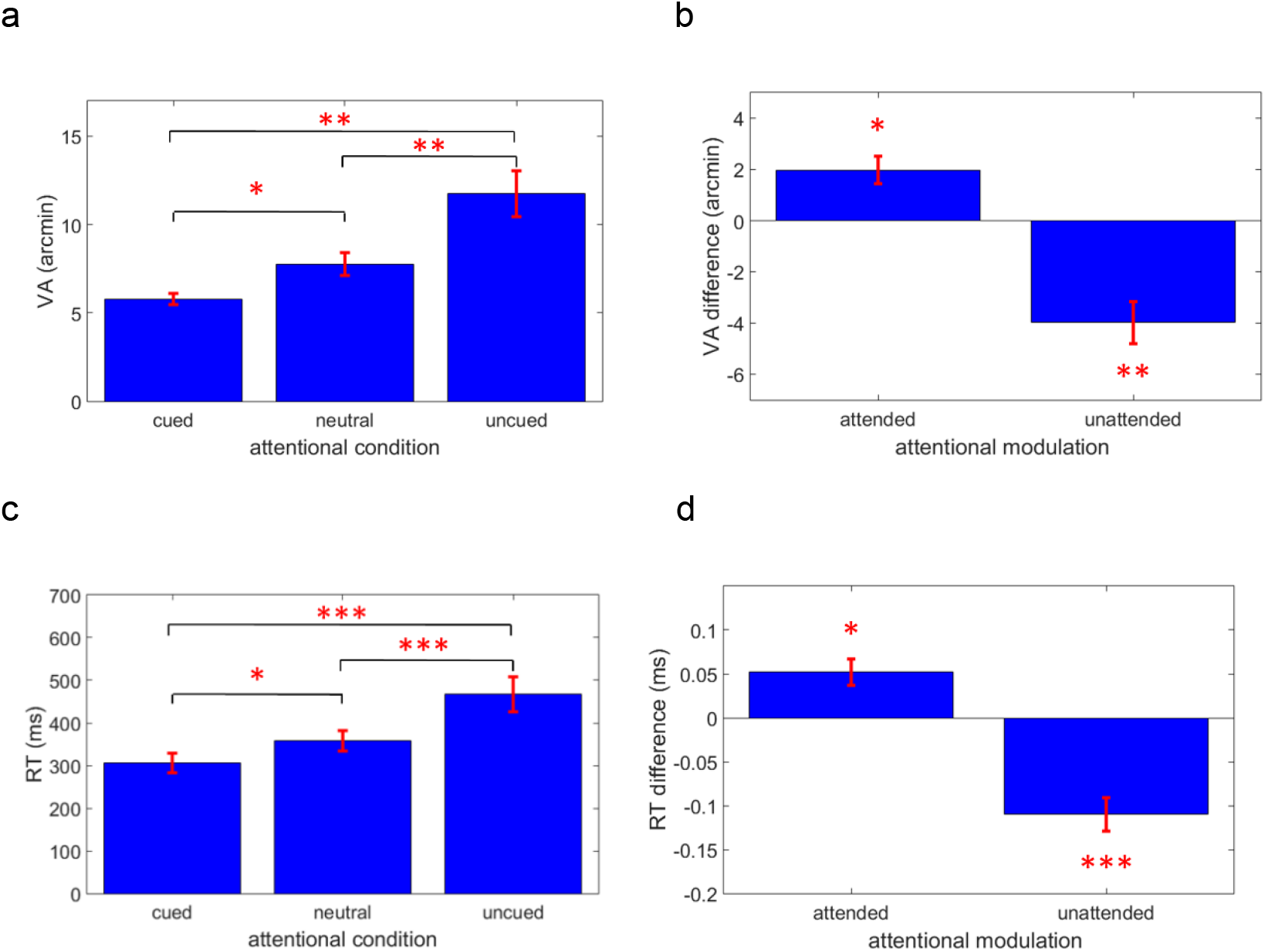
Spatial cueing. (**a**) Mean visual acuity (VA) for the three cue conditions. (**b**) Changes in VA at the attended and unattended locations using the neutral condition as a baseline (VA_neutral_–VA_cued_ and VA_neutral_–VA_uncued_, respectively). (**c**) Mean response time (RT) for the three cue conditions. (**d**) Changes in RT at the attended and unattended locations using the neutral condition as a baseline (RT_neutral_–RT_cued_ and RT_neutral_–RT_uncued_, respectively). In this and following figures, error bars represent one standard error of the mean.

### Effect of spatial cueing on clarity

We replotted in Fig. 3 the spatial cueing data for both clear and blurred targets showing that, under both conditions, VA and RT increased under the cued condition (Bonferroni post-hoc test VA: blurred target: mean difference = 1.528, p=0.02; clear target: mean difference = 2.433, p=0.017) and decreased under the uncued condition (Bonferroni post-hoc test VA: blurred target: mean difference = 3.855, p=0.002; clear target: mean difference = 4.109, p=0.003), when compared with the neutral cue condition. Nevertheless, visual blur significantly influenced VA (three-way RANOVA: F(1,10)=54.203, p<0.001, *η*^2^ =0.844). Notably, a significant interaction was found between visual blur and spatial cueing for VA (three-way RANOVA VA: F(2,20)=3.586, p=0.047, *η*^2^=0.264), but not for RT (3-way RANOVA: F(2,20)=0.142, p=0.869, *η*^2^=0.014). A significant impact of blur (Fig. 3e, VA_blur_-VA_clear_) was observed for both cued and neutral conditions, but not for the uncued condition (Bonferroni post-hoc test VA: cued: mean difference = 1.857 arcmin, p<0.001; neutral: mean difference= 0.952 arcmin, p=0.015; uncued: mean difference= 0.698, arcmin, p=0.074). Most remarkably, the suboptimal attentional conditions (i.e., neutral and uncued) reduced the difference in resolution between blurred and clear stimuli, suggesting that as attention is diverted from the resolution task at hand, visual differences between stimuli having distinct focus are no longer processed.

**Figure 3.**
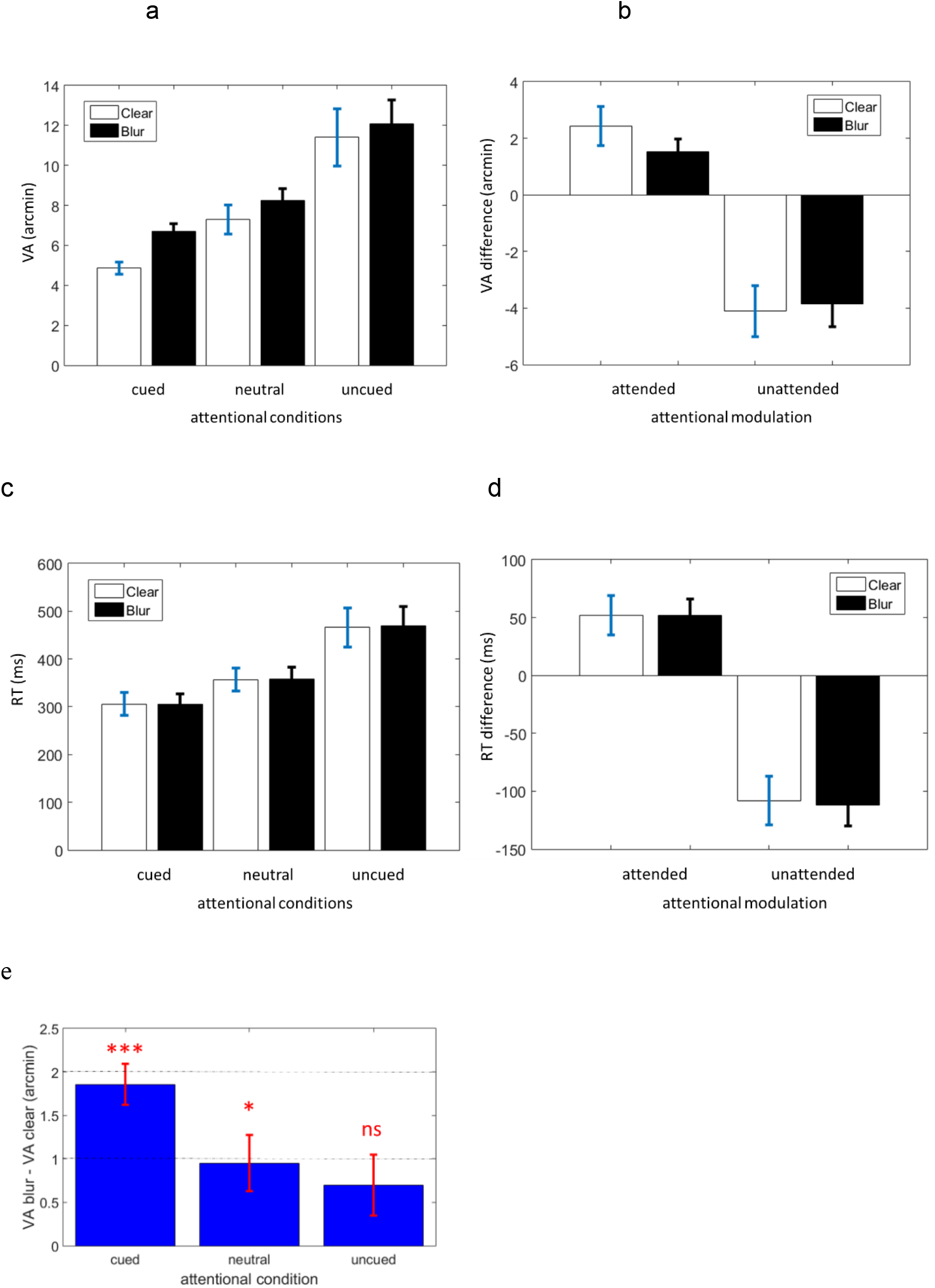
Spatial cueing and visual blur. (**a**) visual acuity (VA) for the three cue conditions and the two blurred conditions. (**b**) Changes in VA at the attended and unattended locations for the two blurred conditions using the neutral condition as a baseline (VA_neutral_–VA_cued_ and VA_neutral_– VA_uncued_, respectively). (**c**) Response time (RT) for the three cue conditions and the two blurred conditions. (**d**) Changes in RT at the attended and unattended locations for the two blurred conditions using the neutral condition as a baseline (RT_neutral_–RT_cued_ and RT_neutral_–RT_uncued_, respectively). (**e**) Effect of visual blur on visual acuity under the three cue conditions. Acuity difference between blurred and clear targets. When attention becomes diverted from the target location, the difference of resolution between blurred and clear images is mitigated.

### Existence of a neuro-optical balance

This equalization of in and out of focus stimulation reveals that the alteration and augmentation of acuity involve an interaction between optical quality of the eye and the movement of attention over the visual field. As depicted in Fig. 4, a paired sampled t-test shows that a defocus system under full attention (attended blur image: 6.72 ± 1.42) exhibits superior performance than a perfectly focused system with reduced attention (unattended clear image: 11.40 ± 5.14); t(21) = 4.58, p<0.001. This indicates that, by increasing a person’s attention, it is possible to compensate for a large drop in optical resolution. Similarly, a defocus system under full attention (attended blur image: 6.716 ± 1.417), has similar performance than a focused system with reduced attention (neutral clear image: 7.292 ± 3.347); as a paired sampled t-test shows no statistical difference t(21) = 1.04, p=0.312. This highlights that for a given acuity level, it is possible to have varying combinations of attention and focus. This highlights that natural modulation of attention (e.g., occurring under prolonged sustained attention [**44-46**]) can be counterbalanced by an increase in optical resolution.

**Figure 4.**
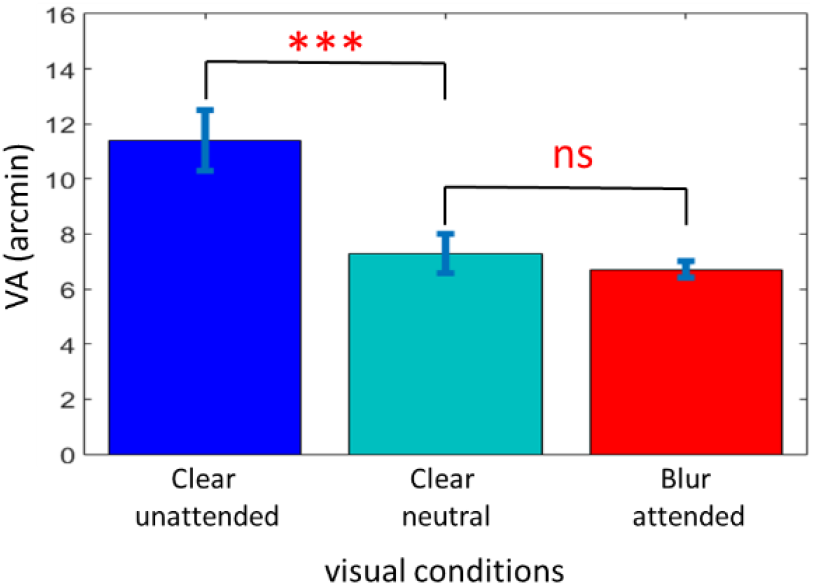
Interaction between attention and blur. Mean VA for different combinations of attention and blur levels. The deployment of attention resulted in a defocused system (blur image) that performs better than, or the same as, a focused system (clear image). This indicates that different combinations of the degree of a person’s attention and his/her optical correction are possible for the visual system to achieve a given acuity resolution.

### Effect of clarity on spatial cueing

Interestingly, the cueing gain (Fig. 5, VA_cued_/VA_neutral_) was significantly augmented in clear targets compared to blurred targets (Paired-samples t-test: mean difference: 0.290, paired t(10)=2.37, p=0.039) whereas the cueing cost (Fig. 5a VA_uncued_/VA_neutral_) did not appear to be influenced by the blurring condition (Paired-samples t-test: mean difference:-0.02, paired t(10)=-0.061, p=0.557). The absence of difference in acuity reduction (from the neutral baseline) for blurred and clear targets is compatible with the idea that, below a certain level of attention (here, referring to the neutral conditions), the information modulated by attention in clear targets becomes similar to that for blurred targets. On the other hand, the total cueing effect was stronger for clear targets than blurred targets, which revealed that spatial attention enhanced acuity more with clear than blurred vision. Albeit speculative, a plausible explanation is that the effective width of the attentional filter is broader for clear vision, because of the augmentation of the suprathreshold components of the image (e.g., towards the higher spatial frequencies) on which attention can operate, as illustrated in Fig. 6.Those findings indicate that the efficiency of attention deployment is somewhat dependent on the image quality of the human eye at a given location. If attention do depend on the retinal blurred image, the particular distribution of blur across the visual field, which is known to vary with refractive populations, could matter in the distribution of spatial attention. This asks whether the retinal blurs resulting from the eye growth are merely an outcome of biological constraints, or could involve some neural feedback mechanisms involved in the regulation of the attentional resources?

**Figure 5.**
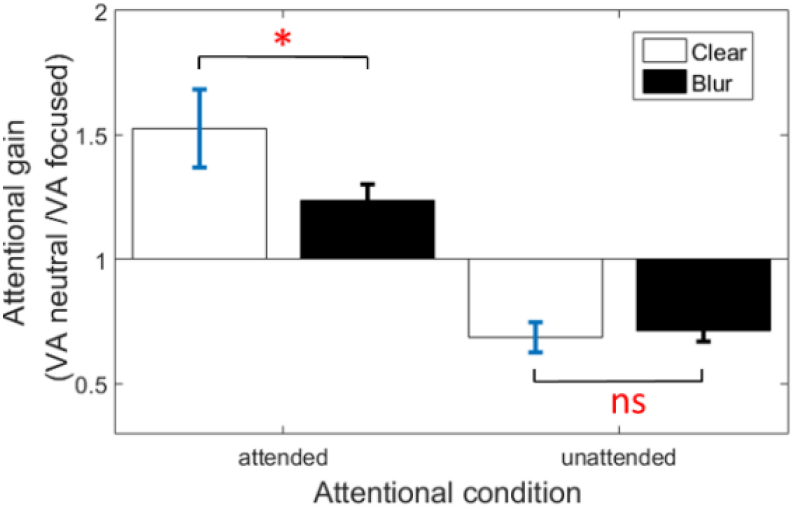
Impact of visual blur on the beneficial effect of attention. Effect of attentional modulation on VA. Ratio in VA between neutral and focused conditions (VA_neutral_/VA_focused_) at cued and uncued locations (i.e., cued and uncued conditions, respectively). Visual blur decreased both beneficial and cost effects of cued attentional orienting, which resulted in a narrower range of resolvable stimulus by the focus of attention.

**Figure 6.**
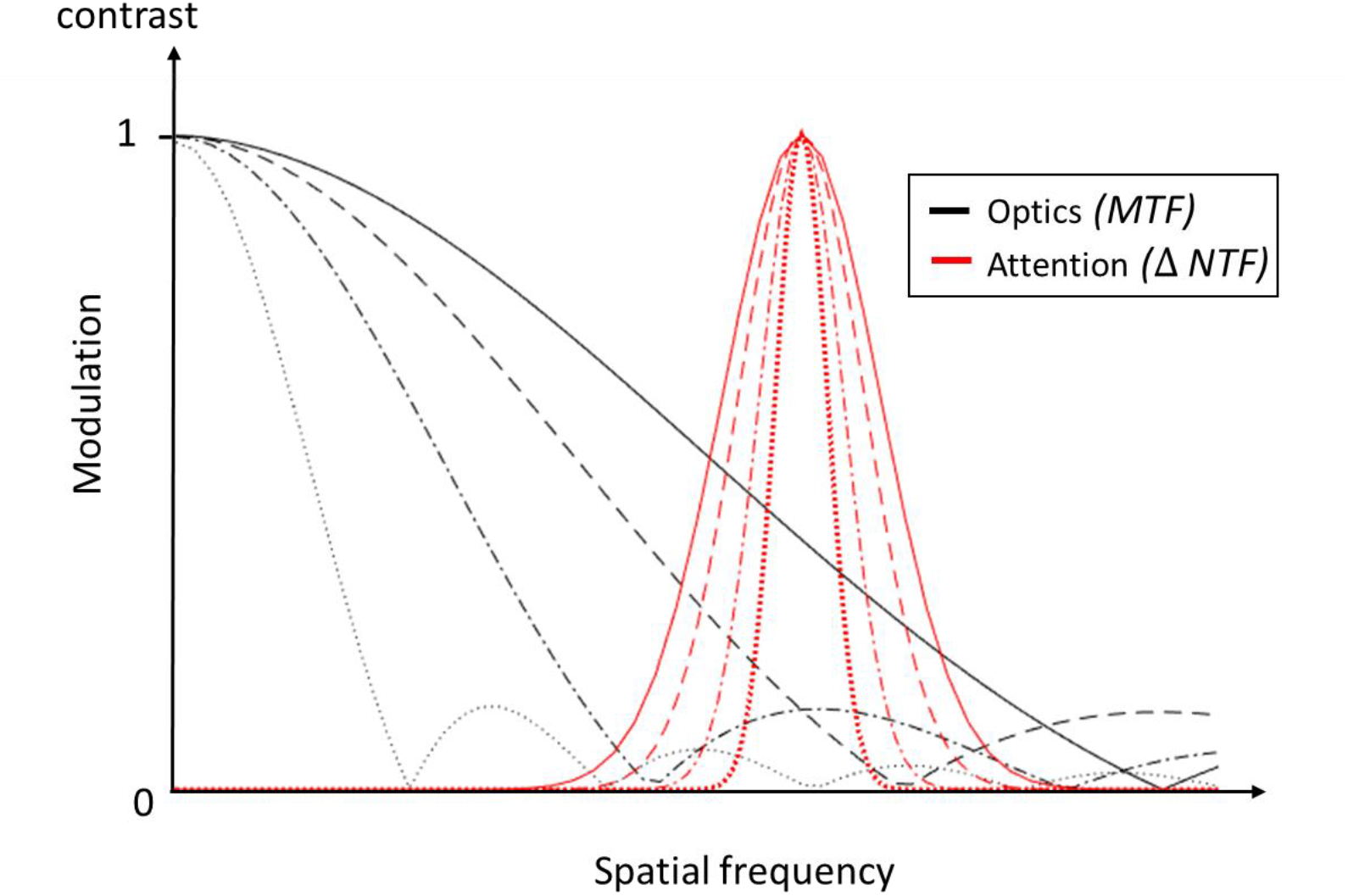
Changes of optical and neural filter. Schematic diagram showing how the gain of a spatially localized attentional filter (ΔNTF) could vary in response to the variation of the modulation transfer function (*MTF*) of the eye for various level of defocus. The rate of the dash line indicates the level of blur associated with the optical and attentional filter. As the magnitude of blur increases, the maximum effective width of the attentional filter is reduced in the highest spatial frequency of the image due to the shrinkage of the area under the modulation transfer function.

### Simulated refractive gain

The data were replotted in terms of visual acuity gain (VA_blur_/VA_clear_) as shown in Fig. 7a. A significant decrease (one-way RANOVA, F(2,20)=7.697, p=0.003, *η*^2^=0.435) in refractive gain by a factor of approximately 1.26 and 1.19 was found under neutral and uncued conditions, respectively, compared to the cued condition, indicating that a certain degree of attention is required for attaining a benefit of sufficient value from blur correction. Fig. 7b shows the simulated refractive gain variations associated with modulation in attention from the neutral condition. It is worth noting that, under suboptimal attentional conditions, the correction of retinal blur may not increase acuity although stimuli are still detected. This finding identifies attention as a prerequisite of vision enhancement in the perifovea.

**Figure 7.**
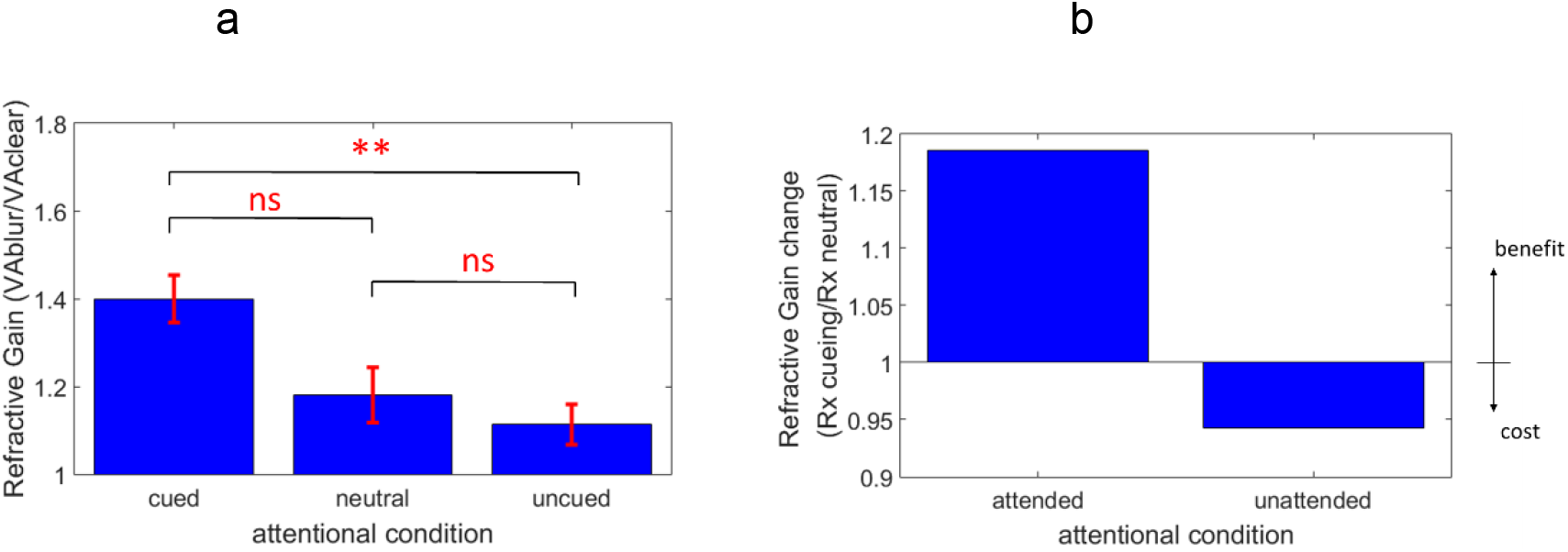
Impact of attention on refractive gain. (**a**) Ratio in VA between blur and clear images (*R*_*x*_=VA_blur_/VA_clear_) as a function of cueing conditions. As attention diminished, the beneficial effect of blur correction decreased. (**b**) Refractive gain change (*R*_*x*_cueing/*R*_*x*_ neutral) from the baseline condition (i.e., neutral attention) associated with attention modulation. Attention focusing resulted in a large increase in the expected gain of blur correction with respect to neutral condition, but only slight reduction accompanied attentional diversion.

## Discussion

In this study, it was demonstrated that acuity depends on both attentional and optical factors, throwing light, for the first time, on the way resolution acuity is modulated by variations in endogenous covert attention and the eye’s focus (Fig. 2-4, 6-7). This complements a large number of studies in the literature that have till now discounted the interaction between attention and optics, focusing only on the separate effects of attention [**18-32**] and ocular optics [**34-40**] per se in acuity resolution.

### Tolerance to retinal blurs

We showed that processing of blur is strongly influenced by the level of attention, and so not simply automatically processed [**10**]. Specifically, it was determined that attention not only improved images that are sharp [**18-27**], but can also drastically improved the resolution of degraded retinal images (Fig. 2a-c) to provide more tolerance to blur. This tolerance suggests the possibility of an adaptive compensation by attention of the retinal blurs degrading vision, which, might enable the eye to rely less on the optical ocular quality. This may be particularly useful to adapt the unwanted spatiotemporal blurs of the visual field that are caused by the motion of the eye and visual targets. Besides, it could allow relaxation of the constraint on optical quality required to achieve a certain performance when recomposing these blurs (as is the trend with progressive addition lenses). On the other hand, under diminished attention (Fig. 3e), an optical enhancement could present the sure advantage to providing greater acuity compared to a blurred image. A riveting question remains how optical modulations affect our attentional responses and resources.

### Attentional limit to supervision

Our results reveal that blurring does modulate the impact of attention. We show that attention boosts more acuity resolution in clear stimuli than blurred stimuli (Fig. 5). While this seems to accord with the idea that attention enhance more retinal stimuli having the highest level of details [**30**], it is also plausible that the increased contrast across spatial frequencies in clear stimuli favor attentionnal modulation. Supposing that clear retinal stimuli are more attentionally demanding, ocular blurs brought by the eye growth could be a way for the visual system to regulate the deployment of attentional resources across the visual field, and adjust the cues of the environment. A practical, and potentially interesting, consequence of a differential attentional modulation is that the decrease of attention can alter the resolution gain of optical correction (Fig. 7): the conditionality of beneficial effects of correction in daily activities can have certain implications for visual technologies aimed at augmenting visual resolution. For instance, a super-resolved optical system [**40**] may only be effective if subjects allocate sufficient attentional resources to a given location. Such contingency of performance requires an individual’s attentional pattern to be considered when determining the level of correction. Indeed, the observed effect of attention and blur and their interaction could vary between individuals because of several factors, including ocular aberration, neural sampling, neural adaptation but also maybe individual attentionnal patterns.

### Dynamic focus correction

An expected, but seminal, finding of this study is that a given level of acuity can involve different combinations of attention and focus (Fig. 4)-that is, a focused system with reduced attention can perform closely to a defocused system under full attention. This provides evidence that changes in attention might be balanced by external or/and changes in optical focus, and vice-versa. Given the incessant modulation of attention with time [**15**], environmental settings [**12**], but also among individuals [**47**], the use of adaptive optical technology could constitute a unique opportunity to adjust the modulation of attention at the optical level. While such optical compensation is not feasible using standard static corrections, given their fixed focus, several emerging technologies, such as spectacles-free display [**1**] and adaptive optics spectacles [**2**], show the potential to dynamically adjust the level of acuity to the movement of attention via an adjustable focus, which could thus timely control the resolution of the neural images. While a dynamic optical correction might be simply based on the person’s feedback, in the case of an automatic optical focus, it would require the ability – not without challenges-to continuously sense and decode the eye responses to access the temporal variations of attention. Combining real-time eye-tracking and control of the optical focus of neural images may open up unique and exciting horizons to respond to individual visual needs, which hold promises for the development of visual aids capable of dynamically linking optical inputs and neural outputs.

### Future development

This study has some limitations. First, the control of ocular aberrations in the extrafoveal regions of the retina was not possible, as, to date, there is no visual simulator capable to correct aberration over a wide viewing angle. Visual simulator correcting the aberrations of the eye are restricted by the isoplanatic patch of the eye to a small visual angle (of about 1-2degrees) that allows testing only one peripheral location at once [**48**]. This severely limits the manipulation of spatial attention across distant locations of the visual field. By using an adaptive optical system incorporating several deformable mirrors and wavefront sensors, it may be possible in the future to enlarge the angular extent of adaptive ocular correction of the eye [**49**], and so the limitation of conventional visual simulators, which incorporate only a single deformable mirror. The use of a wide field AO would help, not just to simulate different retinal blurs, but also compensate for the aberrations of the eye. Such compensation is essential to investigate a possible effect of neural adaptation to natural peripheral blurs in spatial attention. For example, sensitivity to the simulated blurred images could be influenced by individual differences in natural peripheral blurs the eye may be adapted to [**50**]. Nevertheless, little is known about the degree of adaptation to *natural* peripheral aberrations, though recent studies in Yoon Lab suggests that adaptation to defocus could differ with individual refractive errors in myopes and emmetropes [**51**]. Further works will be required to elucidate this. A second limitation of the study is that the interleaved test was restricted to a limited number of retinal conditions. It is plausible that the joint effect of attention and blur may vary as the characteristic of ocular blur changes across the visual field, but a comprehensive understanding of these parameters could involve controlling for the variation of ocular blur across eccentricities, as recently performed in our Lab via a multiscale visual simulator [**52**]. To avoid the confounding effects of the individual ocular aberrations on the simulated retinal blurred images, a sufficiently large blur, producible on the display, was considered, excluding blur conditions with very small and large amounts of retinal blurs. Other limitations could involve the complexity of the stimulus and task. For instance, we show in a recent study [**53**] that, when individual s perform a simple detection task, the effect of exogenous spatial attention on simulated blurred images is small, suggesting a plausible decrease of interaction between blur and attention as the attentional demand required by the task diminishes. Given that exogenous and endogenous attentional filters involve different neuronal pathways, the dynamic of attention might influence an interaction between blur and attention. Further works will be needed to elucidate how these factors affect the processing of blur processing by attention.

## Conclusion

In sum, our results proved, for the first time, a joint effect of optical and attentional factors on acuity resolution, showing that both attention and the eye’s optics matters in the way we perceive things: both the degree of attention and optical correction vary the acuity resolution of the visual system, which potentially allows distinct (neuro-optical) combinations for achieving a given visual resolution. The interaction between those two visual factors is, however, more complex than thought, as acuity enhancement is not equal for focused and defocused images, the movements of attention could modulate an optical correction of the eye. This suggests an important avenue of exploration for adaptive optical technologies. Indeed, the neuro-optical interaction investigated in the present study for localized, defocused images is one only tiny facet of the iceberg, with a more important question perhaps, that is: how, and to which extent, a person’s attention interact with the patterns of blurring on the retina? We believe that further research utilizing new optical advances to control ocular blurs and attention over the entire visual field could contribute to unfolding this mystery. Future works shall examine how the interaction that may link the brain and the optics of the eye spreads across the visual field in real-world contexts.

## Methods

### Experimental design

Subjects were asked to fixate on a small cross displayed at the center of the monitor screen (Fig. 1, “+”; size, 0.5° x 0.5°). The endogenous covert attention of the subject was manipulated using a central cue preceding the visual stimuli. On cued and uncued trials, a central cue was displayed (size, 18 x 0.6 arcmin; exposure time, 293 ms), pointing at either one of the two upcoming Landolt C locations, to which the observer was required to allocate attention. In neutral trials, two central cues were displayed, pointing at both upcoming Landolt C locations. Two Landolt Cs (“target” and “non-target”), with different gap orientations, were then presented simultaneously after a 300 ms inter-stimulus-interval. At the end of the presentation sequence, a central line symbol (size, 18 x 0.6 arcmin) was displayed to point the location of the target Landolt C.

Three cue conditions were tested: (1) in the cued condition, the cue pointed towards the target Landolt C location; (2) in the uncued condition, the cue pointed towards the non-target Landolt C location; and (3) in the neutral condition, two cues were displayed, which pointed towards both target and non-target locations. Each cue condition was displayed for one-third of the response trials in each session (cued: uncued: neutral = 1:1:1). It should be noted that the participants were required to direct their attention as instructed by the cue(s), and that they were not informed of the proportions of the three cue conditions.

The experimental procedures were approved by the University Committee for the Protection of Human Subjects (HSEARS20170103003), and the research was conducted according to the principles expressed in the Declaration of Helsinki. Informed consent was obtained from each participant. All those involved, except two of the observers, the authors J.T.L and D.L.E, were inexperienced with psychophysical procedures and not informed of the purpose of the experiments.

### Simulated defocus blur

Visual performance was assessed under both clear (zero blur) and blurred (defocus blur, 2 waves of RMS wavefront error, about 1.25D) conditions. Note that the blurred Landolt C stimuli were graphically generated by convolution of the two-dimensional luminance profile of the Landolt C and a point spread function, *h*(*x, y*), of a 5 mm pupil (2*r*):

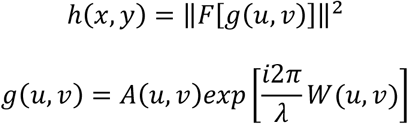

where *F* denotes Fourier transform, *A*(*u, v*) denotes pupil function, *W*(*u, v*) denotes wavefront aberration described by a set of Zernike polynomials [**54**]. In this study, the visual stimuli (i.e. Landolt C letters) were presented in black against a green background, and therefore λ was set to 550 nm. The wavefront aberration was calculated from the Zernike defocus polynomials 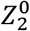, as:

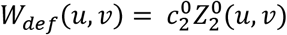

The amount of defocus used in these experiments is given by [**55**]:

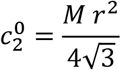

where *M* denotes the spherical equivalent in diopters, *r* the radius of the simulated pupil.

### Visual acuity measurement

Eleven young adults with corrected-to-normal vision (age 24- 39; VA 6/6 or better in each eye) performed a Landolt C acuity task at two locations in the near-peripheral visual field (Fig. 1, 8° eccentricity; quadrant, NW and SE; along 135°/315° with horizontal). The placement of the pair of stimuli at the intercardinal locations aimed to minimize field performance and attentional asymmetries. The visual task was to identify the orientation of a gap in the “target” Landolt C (6AFC: 30°, 90°, 150°, 210°, 270°, 330°). The stroke width was one-fifth of the Landolt C size. The stimulus exposure duration was 33 ms. Viewing was binocular.

An interleaved 1-down-1-up staircase procedure that converged on the 50 % correct level (adjusted for 6AFC, guess rate=1/6) was used to control the gap size and measure the visual acuity threshold. There was a total of 12 interleaved staircases for various stimulus conditions (i.e., three cue conditions x two locations x two blur levels). The acuity test was repeated a total of 15 times for each stimulus condition over five sessions (approximately 5400 response trials in total), and the average value of the threshold measurements was taken as visual acuity. Observer responded using a keyboard and audio feedback was provided after each response. The time taken for an observer to respond after the offset of the Landolt C stimuli was measured as response time. Only response taking place after the response cue offset were considered, response time longer than 1 second were excluded.

A 32-inch Dell LCD monitor (screen resolution, 3840 x 2160; background luminance, 25 cd/ *m*^2^; contrast, 1100 %) was used to display visual stimuli. Viewing distance was 1m. Throughout the experiment, an Eye tracker (Tobbi TX 300) monitored eye fixation, from the onset of the cue to the offset of the target. Trials with unstable fixation (eye movements > 2°) were discarded.

## Notes

### Competing Interest Statement

The authors have declared no competing interest.

